# Differential representations for affective and informative components of reward in the striatum and hippocampus

**DOI:** 10.1101/2024.09.20.614186

**Authors:** Xinxu Shen, Kainan S. Wang, Vishnu P. Murty, Mauricio R. Delgado, David V. Smith

## Abstract

Receipt of a reward is composed of affective and informative components, which are often intertwined in most reward-processing and decision-making tasks. Our previous work allowed us to identify regions that were more strongly engaged for the affective and informative components of reward upon receipt (Smith et al., 2016) and showed that the ventral striatum responds to both affective and informative components of reward processing. However, the limited spatial resolution and coarse analytical approaches made it hard to understand how these different components of reward were represented in the brain, wherein similar engagement may not necessitate similar representations. In our current study, we used high-resolution functional magnetic resonance imaging (voxel size: 1.75 mm^3^) and representational similarity analysis to investigate the representation of affective and informative components of reward receipt in the striatum and the hippocampus, another region that is sensitive to reward information. We found a differential representation of affective and informative components of reward receipt in the striatum and the hippocampus, with no difference in representation between the two structures. However, in the dorsal striatum, we found that representations were stronger for affective rather than the informative components of reward receipt. Finally, we observed that the ventral striatum was sensitive to the predictiveness of information, such that across-run pattern similarity in the ventral striatum increased with the predictiveness of information. In sum, our results provide evidence of how affective and informative components of rewards are represented in the striatum and the hippocampus, potentially indicating a differential coding schema for the dorsal and the ventral striatum.

## 1. Introduction

Receipt of a reward is associated with multiple components, notably affective components that modulate our affect (e.g., promote positive emotions) and informative components that provide information that guides future behavior (Schultz, 2006). To illustrate, consider a gambling scenario where monetary feedback has a dual effect: it instantly influences one’s affect, evoking happiness upon a financial gain, while concurrently furnishing critical insights about the game’s structure, which in turn shapes expectations for future gains. The interplay between these affective and informative components of reward receipt underscores their distinctive roles in optimizing behavior and promoting well-being. Thus, there is a growing interest in investigating whether these two components of reward receipt are differentially represented in the brain, influencing decision-making processes. Previous research has made efforts to examine how the brain processes affective and informative components of reward receipt (Smith et al., 2016), with overlapping engagement of the same region in response to both of these features of reward. Yet, there remains a knowledge gap regarding whether these components are represented similarly in the brain even when a region is equally activated by both, possibly due to limitations in resolution and advanced neuroimaging analysis techniques. Therefore, the current study used high-resolution neuroimaging techniques combined with representational similarity analysis to uncover the representations of affective and informative components of reward receipt in the brain.

Several lines of research have uncovered the role of the striatum, both its ventral and dorsal portions, in response to affective and informative components of reward receipt (Dennison et al., 2022; Smith & Delgado, 2015; Wang et al., 2016). For example, Delgado et al (2000) used a paradigm to isolate the effect of rewards and observed differential activation to the valence of outcomes in the striatum, suggesting an engagement in processing the affective components of reward receipt, which is consistent with animal research showing the linkage between the dorsal striatum and reward-related activity (Hollerman et al., 1998). In addition, previous research also found the engagement of the striatum in the processing of information-related signals. For example, a study found that feedback-related processing in the dorsal striatum is influenced by the amount of information provided by negative feedback during a paired-associate learning task (Tricomi & Fiez, 2012). Together, these studies suggested that the striatum is engaged in the process of both affective and informative components of reward receipt. However, few studies have directly compared affective and informative components of rewards in one study design.

Therefore, to answer the question of whether affective and informative components of reward receipt are represented differently in the brain, we previously developed a task that allowed us to directly compare affective and informative reward components upon reward receipt (Smith et al., 2016). Specifically, participants completed two card guessing tasks: one task asked participants to accrue points through a guessing game (affective card task) while another task asked participants to learn the association between the feedback and their choices (informative card task). Using this design, we found that both affective and informative components of reward receipt evoked activation within the nucleus accumbens, a subregion of the ventral striatum (Smith et al., 2016).

While the focus on both the affective and informative components of reward receipt has mainly been in the striatum, emerging evidence has been highlighting a role in the hippocampus. For example, the hippocampus has been shown to be primarily involved in processing informative components of reward receipt related to learning and memory (Charpentier et al., 2018; Chen et al., 2016). The hippocampus also receives and integrates affective information with other types of information, such as contextual information, and forms memories (Cowan et al., 2021). We, therefore, hypothesize that the hippocampus might be important for the representation of affective and informative components of reward receipt as well. We further hypothesize that there might be dissociable processing of affective and informative components of reward receipt between the striatum and hippocampus, such that the striatum is primarily involved in processing the affective component of reward receipt (Delgado et al., 2000; Smith et al., 2016), whereas the hippocampus is primarily involved in processing the informative component of reward receipt.

One way to resolve whether a given region differentially represents affective and informative information, even when they both activate a given region, is to use the representational similarity analysis. Representational similarity analysis—a multivariate technique—can identify patterns of neural activity for affective and informative components of reward receipt and compare representations in the striatum and the hippocampus (Chavez & Heatherton, 2015; Devereux et al., 2013; Popal et al., 2019), which helps better detect differences across processes. For example, two processes that evoke suprathreshold activation in a region may be tied to distinct patterns (Woo et al., 2014). It is hard to detect these differences using univariate analysis, but representational similarity analysis could help detect differences and similarities that eluded our original work.

We further propose that there might be dissociations within the striatum and the hippocampus in representing the affective and informative components of reward receipt. High-resolution neuroimaging technique is important for detecting the dissociations within a structure. We suggest that it is possible that our prior work was unable to uncover distinct representations of affective and informative components of reward receipt due to blurred anatomical boundaries and poor spatial resolution. For example, the ventral and dorsal striatum might play different roles in processing affective and information components of reward receipt, but it is difficult to detect dissociation without high-resolution neuroimaging techniques.

Therefore, in the current study, we used high-resolution fMRI (Olman & Yacoub, 2011; Zaretskaya, 2021) to identify smaller and more specific regions within the striatum, such as dorsal vs. ventral striatum, and within the hippocampus, such as anterior vs. posterior hippocampus, to better understand the representation of affective and informative components of reward receipt in these regions. Rather than focusing on univariate activation, we used Representational Similarity Analysis (RSA)—to differentiate distinct activation patterns to capture different representations of affective and informative components of reward receipt. Specifically, we adapted the paradigm from our previous work and collected high-resolution fMRI data. We separated affective and informative components of reward receipt in two card games using different instructions to emphasize either the affective or informative components of reward receipt. Our analyses focused on two pre-registered questions (http://aspredicted.org/~N47uU5Qjkt). First, how are affective and informative components of reward receipt represented in the striatum and hippocampus? Second, are the striatum and hippocampus sensitive to affective and informative reward magnitude?

## 2. Materials and Methods

### 2.1. Participants

Twenty-four participants (Age: 18-28, 8M, 16F) were recruited from Rutgers University – Newark to participate in this study. Participants were excluded for fd_mean values greater than 1.5 times the interquartile range, or for tSNR values below the lower bound of 1.5 times the interquartile range, resulting in five exclusions. All participants were screened before data collection to rule out current major psychiatric or neurologic illness, as well as MRI contraindications. All participants gave written informed consent as part of a protocol approved by the Institutional Review Board of Rutgers University -- Newark.

### 2.2. Procedures

#### Affective card task (ACT)

In the affective card game (figure 1A), participants were instructed to earn enough points to play in a bonus game. On each trial, participants were presented with three decks of cards for 2.5 seconds or until selecting a deck. Failure to respond within the required interval resulted in no feedback, which was shown as a black circle. After a card was chosen, a variable fixation interval of 4.5 to 7.5□seconds was presented before feedback from the chosen deck was presented for 1□second on the screen. Feedback was cards depicting a variable number of points (1, 2, 3, 4) with equal probabilities in each deck (1 represented low reward and 4 represented high reward). Trials were separated by a variable intertrial interval of 6 to 12 seconds (ITI).

**Figure 1.**
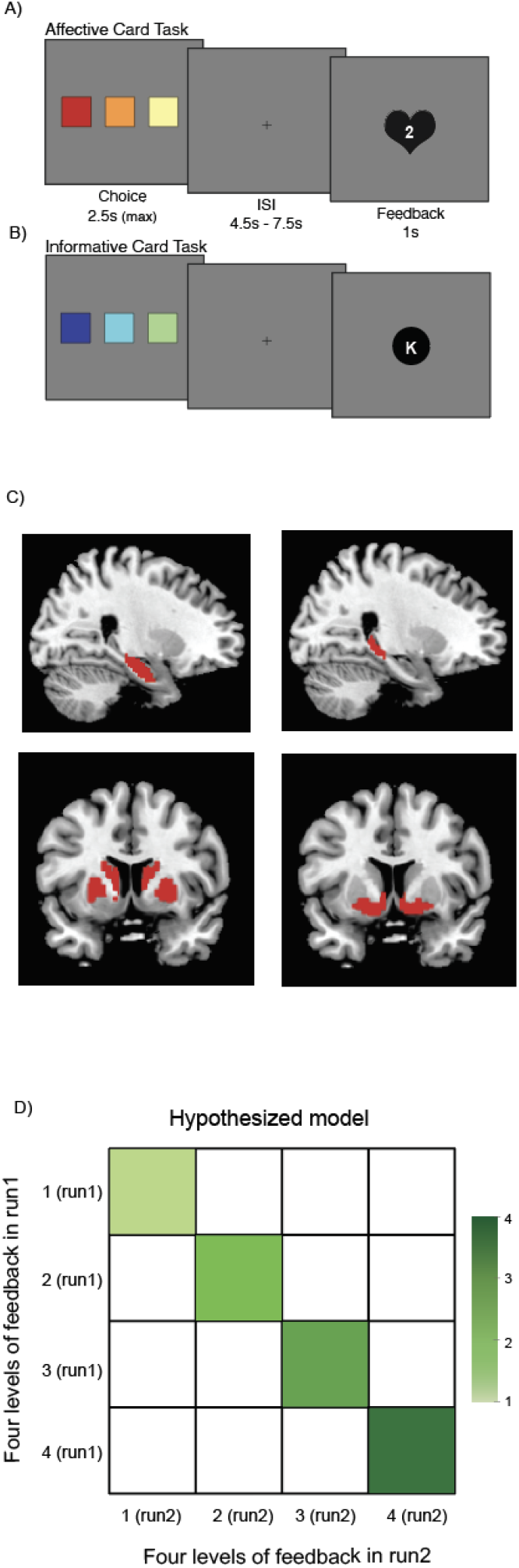
A) Affective card task. Participants were instructed to choose from three card decks and received three levels of feedback (1-4 points). The goal of the task was to earn enough points to play in the bonus game. One point represented low reward/affect and four points represented high reward/affect. B) Informative card task. Participants received four different letter feedback (D,K,X,Z). The goal of the task was to learn the contents of each deck of cards, which will be used in the bonus game. Each letter appeared with different probabilities in each card deck. C) Four regions of interest: anterior hippocampus, posterior hippocampus, dorsal striatum, and ventral striatum. D) A hypothesized model that assumed across-run pattern similarity increased linearly as reward magnitude increased.

#### Informative card task (ICT)

In the informative card task (figure 1B), the goal for the participants was to learn the contents of each deck of cards because the bonus game would explicitly test this knowledge. The task was the same as the affective card task, except that feedback was cards depicting variables in four different letters (D, K, X, Z) with different probabilities in each deck. The point distributions within each deck were as follows (10% represented rare information and 40% represented more common and predictable information):

Deck 1: D (40%), K (30%), X (20%), Z (10%).

Deck 2: D (30%), K (20%), X (10%), Z (40%).

Deck 3: D (20%), K (10%), X (40%), Z (30%).

Different from the objective probability of feedback, we designed this task such that subjective probabilities experienced by the participant changed based on accumulated evidence. Subjective probability represents acquired and updated information and differs across participants. We assumed that participants started both tasks with flat expectations that were updated on a trial-to-trial basis. We calculated subjective probability for each feedback type in the section below and we used subjective probability, instead of objective probability, in our analysis because subjective probability reflects learning and processing of informative components of reward receipt. Participants completed two runs of affective card task (ACT) and two runs of informative card task (ICT). Each run consisted of 45 trials. These runs were interleaved, and order was counterbalanced across participants.

#### Bonus game

In each trial of the bonus game (45 trials per participant), participants were presented with a letter followed by two decks of cards and asked to choose the deck that was most likely to contain the letter. Thus, knowing the complete composition of each deck from the information card task is important to answer the Bonus Game questions correctly. There was no time limit for the bonus game. Participants received their compensation after they completed the bonus game.

### 2.3. Behavioral analyses

In order to determine the subjective value for informative reward components of reward receipt, we categorized each unique combination of card choice and feedback as a distinct instance. For the first run, the frequency of each instance was tracked, starting at one and incrementing by one with each subsequent occurrence (e.g., the second instance of a participant selecting card one and receiving feedback X would result in an updated frequency of two). The total frequency of each instance was then divided into four equal quintiles, with the least frequent feedback assigned to the first quintile and the most frequent to the fourth quintile. Because the second run was influenced by prior knowledge from the first run, the frequency for instances in the first quintile started at one, whereas the frequency for instances in the second quintile began at two, and so forth. Finally, a win-stay-lose-shift model was executed to calculate the probability of participants selecting the same card deck for each feedback type, thereby allowing us to examine the effect of different feedback types on participants’ choice behavior. All statistical analyses were conducted in Python version 3.10.12. ANOVA models were run using the “statsmodels” package to investigate differences in the probability of choosing the same card across different levels of feedback.

### 2.4. Neuroimaging data acquisition

Neuroimaging data were collected at the Rutgers University Brain Imaging Center (RUBIC) using a 3.0 Tesla Siemens Prisma scanner equipped with a 12-channel head coil (repetition time = 2 seconds). Functional images sensitive to blood-oxygenation-level-dependent (BOLD) contrast were acquired using a single-shot T2*-weighted echo-planar imaging sequence with slices parallel to the axial plane [GRAPPA with R = 2; repetition time (TR): 2000 ms; echo time (TE): 28 ms; matrix 128 × 128; field of view (FOV): 204 mm; voxel size 1.75 × 1.75 mm; 30 slices (10% gap; flip angle: 90 degrees). High-resolution structural scans covering the whole brain (TR: 1900 ms; TE: 2.52 ms; matrix 256 × 256; FOV: 256 mm; voxel size 1.0 × 1.0 × 1.0 mm; 176 slices; flip angle: 9 degrees) were acquired to facilitate coregistration and normalization of functional data.

### 2.5. Preprocessing of neuroimaging data

Neuroimaging data were converted to the Brain Imaing Data Structure (BIDS) using HeuDiConv version 0.5.4 (Halchenko et al., 2019). Results included in this manuscript come from preprocessing performed using *fMRIPrep* 20.1.0 (Esteban et al., 2018b, 2018a). The details described below are adapted from the *fMRIPrep* preprocessing details; extraneous details were omitted for clarity.

#### 2.5.1. Anatomical data preprocessing

The T1-weighted (T1w) image was corrected for intensity non-uniformity (INU) with N4BiasFieldCorrection (Tustison et al. 2010), distributed with ANTs 2.2.0 (Avants et al. 2008, RRID:SCR_004757), and used as T1w-reference throughout the workflow. The T1w-reference was then skull-stripped with a Nipype implementation of the antsBrainExtraction.sh workflow (from ANTs), using OASIS30ANTs as target template. Brain tissue segmentation of cerebrospinal fluid (CSF), white-matter (WM) and gray-matter (GM) was performed on the brain-extracted T1w using fast (FSL 5.0.9, RRID:SCR_002823, Zhang, Brady, and Smith 2001). Volume-based spatial normalization to two standard spaces (MNI152NLin2009cAsym, MNI152NLin6Asym) was performed through nonlinear registration with antsRegistration (ANTs 2.2.0), using brain-extracted versions of both T1w reference and the T1w template. The following templates were selected for spatial normalization: ICBM 152 Nonlinear Asymmetrical template version 2009c [Fonov et al. (2009), RRID:SCR_008796; TemplateFlow ID: MNI152NLin2009cAsym], FSL’s MNI ICBM 152 non-linear 6th Generation Asymmetric Average Brain Stereotaxic Registration Model [Evans et al. (2012), RRID:SCR_002823; TemplateFlow ID: MNI152NLin6Asym],

#### 2.5.2. Functional data preprocessing

For each of the 4 BOLD runs found per subject (across all tasks and sessions), the following preprocessing was performed. First, a reference volume and its skull-stripped version were generated using a custom methodology of fMRIPrep. Head-motion parameters with respect to the BOLD reference (transformation matrices, and six corresponding rotation and translation parameters) are estimated before any spatiotemporal filtering using mcflirt (FSL 5.0.9, Jenkinson et al. 2002). BOLD runs were slice-time corrected using 3dTshift from AFNI 20160207 (Cox and Hyde 1997, RRID:SCR_005927).

The BOLD reference was then co-registered to the T1w reference using flirt (FSL 5.0.9, Jenkinson and Smith 2001) with the boundary-based registration (Greve and Fischl 2009) cost-function. Co-registration was configured with nine degrees of freedom to account for distortions remaining in the BOLD reference. The BOLD time-series (including slice-timing correction when applied) were resampled onto their original, native space by applying the transforms to correct for head-motion. These resampled BOLD time-series will be referred to as preprocessed BOLD in original space, or just preprocessed BOLD.

The BOLD time-series were resampled into standard space, generating a preprocessed BOLD run in MNI152NLin2009cAsym space. First, a reference volume and its skull-stripped version were generated using a custom methodology of fMRIPrep. Corresponding “non-aggressively” denoised runs were produced after such smoothing. Additionally, the “aggressive” noise-regressors were collected and placed in the corresponding confounds file. Several confounding time-series were calculated based on the preprocessed BOLD: framewise displacement (FD), DVARS and three region-wise global signals. FD was computed using two formulations following Power (absolute sum of relative motions, Power et al. (2014)) and Jenkinson (relative root mean square displacement between affines, Jenkinson et al. (2002)). FD and DVARS are calculated for each functional run, both using their implementations in Nipype (following the definitions by Power et al. 2014).

The three global signals are extracted within the CSF, the WM, and the whole-brain masks. Additionally, a set of physiological regressors were extracted to allow for component-based noise correction (CompCor, Behzadi et al. 2007). Principal components are estimated after high-pass filtering the preprocessed BOLD time-series (using a discrete cosine filter with 128s cut-off) for anatomical (aCompCor). This subcortical mask is obtained by heavily eroding the brain mask, which ensures it does not include cortical GM regions.

For aCompCor, components are calculated within the intersection of the aforementioned mask and the union of CSF and WM masks calculated in T1w space, after their projection to the native space of each functional run (using the inverse BOLD-to-T1w transformation). Components are also calculated separately within the WM and CSF masks. For each CompCor decomposition, the k components with the largest singular values are retained, such that the retained components’ time series are sufficient to explain 50 percent of variance across the nuisance mask (CSF, WM, combined, or temporal). The remaining components are dropped from consideration. The head-motion estimates calculated in the correction step were also placed within the corresponding confounds file.

All resamplings were performed with a single interpolation step by composing all the pertinent transformations (i.e., head-motion transform matrices, susceptibility distortion correction when available, and co-registrations to anatomical and output spaces). Gridded (volumetric) resamplings were performed using antsApplyTransforms (ANTs), configured with Lanczos interpolation to minimize the smoothing effects of other kernels (Lanczos 1964).

### 2.6. Additional Preprocessing and Construction of Regions of Interest (ROIs)

After running FMRIprep, we did brain extraction in FSL using FEAT version 6.0. Our primary hypotheses concerned a set of four anatomic regions: anterior hippocampus, posterior hippocampus, dorsal striatum (caudate/putamen), and ventral striatum. Rather than defining functional ROIs via a whole-brain searchlight, we defined ROIs from Harvard-Oxford atlas. We divided the hippocampus lengthwise into equal segments delineating the anterior and posterior hippocampus (y-coordinate = -27). We used probabilistic connectivity striatal atlas to derive our dorsal and ventral striatum masks. We used the executive sub-region as the dorsal striatum mask and the limbic sub-region as the ventral striatum mask.

### 2.7. Neuroimaging analyses

Neuroimaging analyses used FSL version 6.0.4. We specifically focused on the activation analyses to investigate how affective and informative components of reward receipt were associated with BOLD responses. The activation model utilized a general linear model (GLM) with local autocorrelation correction using FILM pre-whitening. Our model focused on the brain activation evoked during the feedback phase of the affective card task and the informative card task and used a total of six task regressors. Four regressors of interest included four levels of feedback for either the affective card task (ACT) or the informative card task (ICT) (duration = 750 ms). Two regressors of no interest included the decision phase and missed trials (i.e., failures to respond; duration = 2000 ms). The activation model included additional regressors of no interest that controlled for six motion parameters (rotations and translations), the first six aCompCor components explaining the most variance, non-steady state volumes, and the framewise displacement (FD) across time. Finally, high-pass filtering (128s cut-off) was achieved using a set of discrete cosine basis functions. We combined data across runs, for each participant, using a fixed-effects model.

### 2.8. Representational similarity analysis

The representational similarity analysis (RSA) framework consists of modeling the observed similarity structure of activation patterns with our specified model similarity structures. We followed standard procedures for using RSA, which was done using the nltools in python (dartbrains.org/content/RSA.html). Specifically, for a given subject and an ROI, task-related beta coefficients for each of the 8 conditions of interest (four levels of affective feedback and four levels of informative feedback) were assembled into a condition-by-voxel activity pattern. We then computed the pattern similarity across each beta image, which generated an associated distance matrix within each ROI. Next, we extracted mean within- and across-task similarity for the affective card task and the informative card task. Paired t-tests and ANOVAs were performed to examine whether affective and informative components of reward receipt were represented differently in the striatum and hippocampus. Finally, we constructed a hypothesized model (figure 1D) that assumed that brain activation was sensitive to reward feedback magnitude. We then correlated our hypothesized model with the observed similarity structure of activation patterns to investigate whether the striatum and hippocampus are sensitive to affective and informative reward magnitude.

## 3. Results

### 3.1. Behavioral results

We analyzed the behavioral data to confirm the separation of affective and informative components of reward receipt. We did not find any difference in the probability of choosing the same card across the four affective feedback types (*p* = 0.82) (figure 2A), indicating that the points received in the affective card task did not provide information that guided choices. Different from the informative card task, we found that in the informative card task, the probability of choosing the same card increased with how common the feedback was numerically (figure 2B). In other words, participants were more likely to choose the same card if there was no surprise with the informative feedback. An analysis of variance (ANOVA) on the probability of choosing the same card yielded significant variation among the four levels of informative feedback (F = 12.38, *p* < .001). A post hoc Tukey test showed that the probability of choosing the same card after receiving informative feedback from the first quintile (more salient information) was significantly lower than the probability after receiving feedback from the second quintile (*p* = 0.01), the third quintile (*p* = 0.03), and the fourth quintile (more predictive information) (*p* = 0.03).

**Figure 2.**
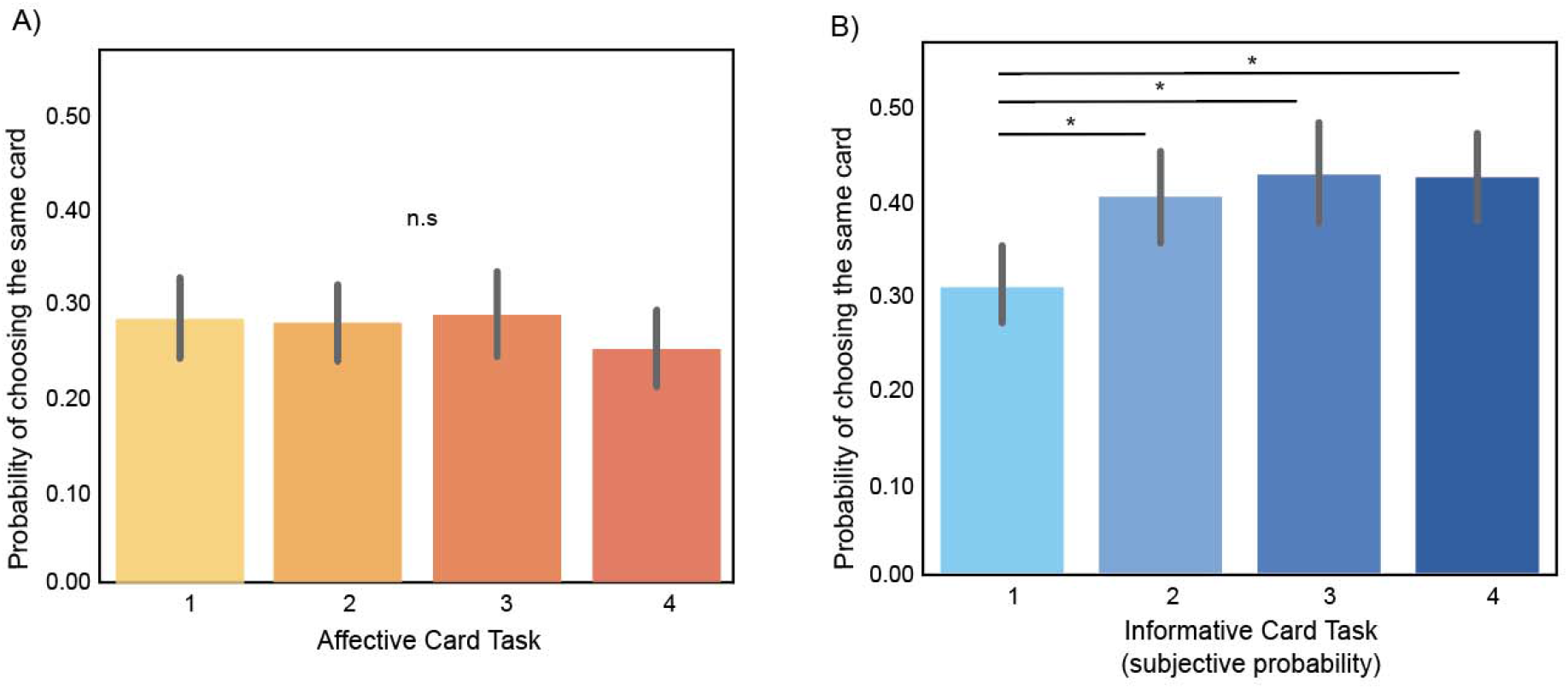
A) The probability of choosing the same card across four levels of affective feedback. There was no relationship between card choice and levels of feedback in the affective card task. B) The probability of choosing the same card across the four levels of informative feedback. The probability of choosing the same card increased with the subjective probability of informative feedback. Participants were more likely to stick to previous card choices when they saw common feedback than salient feedback.

### 3.2. Representation of affective and informative components of reward receipt in the striatum and the hippocampus

Next, we examined how affective and informative components of reward receipt are represented in the striatum and the hippocampus. We conducted the region of interest (ROI) analyses using the dorsal striatum, ventral striatum, anterior hippocampus, and posterior hippocampus, as a priori regions of interest (see pre-registration) (figure 1C). We found that, in each ROI, there was significantly higher within-task versus across-task pattern similarity for both the affective and informative card tasks (ACT: dorsal striatum: t(18) = 11.2, *p* < 0.01; ventral striatum: t(18) = 24.3, *p* < 0.01; anterior hippocampus: t(18) = 55.9, *p* < 0.01; posterior hippocampus: t(18) = 21.7, *p* < 0.01) (ICT: dorsal striatum: t(18) = 7.0, *p* < 0.01; ventral striatum: t(18) = 21.3, *p* < 0.01; anterior hippocampus: t(18) = 24.5, *p* < 0.01; posterior hippocampus: t(18) = 16.7, p < 0.01), suggesting that the striatum and the hippocampus exhibit distinct neural patterns when engaged in different tasks. High within-task pattern similarity implies that the neural representations were more consistent within the same task and lowered across-task pattern similarity implies that the neural patterns were less similar between different tasks. Together, these results suggested that both the striatum and the hippocampus differentially represent affective and informative components of reward receipt.

Then, we investigated whether affective and informative components of reward receipt were represented more strongly in each ROI. We compared within-task pattern similarity for affective card tasks to within-task pattern similarity for informative card tasks. We found that affective components of reward receipt were more strongly represented than informative components of reward receipt in the dorsal striatum (t(36) = 3.22, p = 0.003), such that within-task pattern similarity for the affective components of reward receipt was significantly higher than within-task pattern similarity for the informative component of reward receipt in the dorsal striatum (figure 3C). There was no difference in pattern similarity between affective and informative components of reward receipt in the other three a priori ROIs (figure 3A,B,D) (anterior hippocampus: *p* = 0.79; posterior hippocampus: *p* = 0.16; ventral striatum: *p* = 0.55). To determine if the differences in the dorsal striatum were significantly greater than the other ROIs, we ran an ANOVA but found no main effect of ROIs (F(1,18) = 0.20, *p* = 0.66) or main effects of the tasks (ACT vs. ICT) (F(1,18) = 3.2, *p* = 0.09) and there was no interaction between ROIs and tasks (F(1,18) = 2.24, *p* = 0.15). Together, our findings suggest differential representation of affective and informative components of reward receipt in the hippocampus and the striatum, with no difference in representation between the hippocampus and the striatum. However, the dorsal striatum represented affective components of reward receipt to a greater extent than informative components.

**Figure 3.**
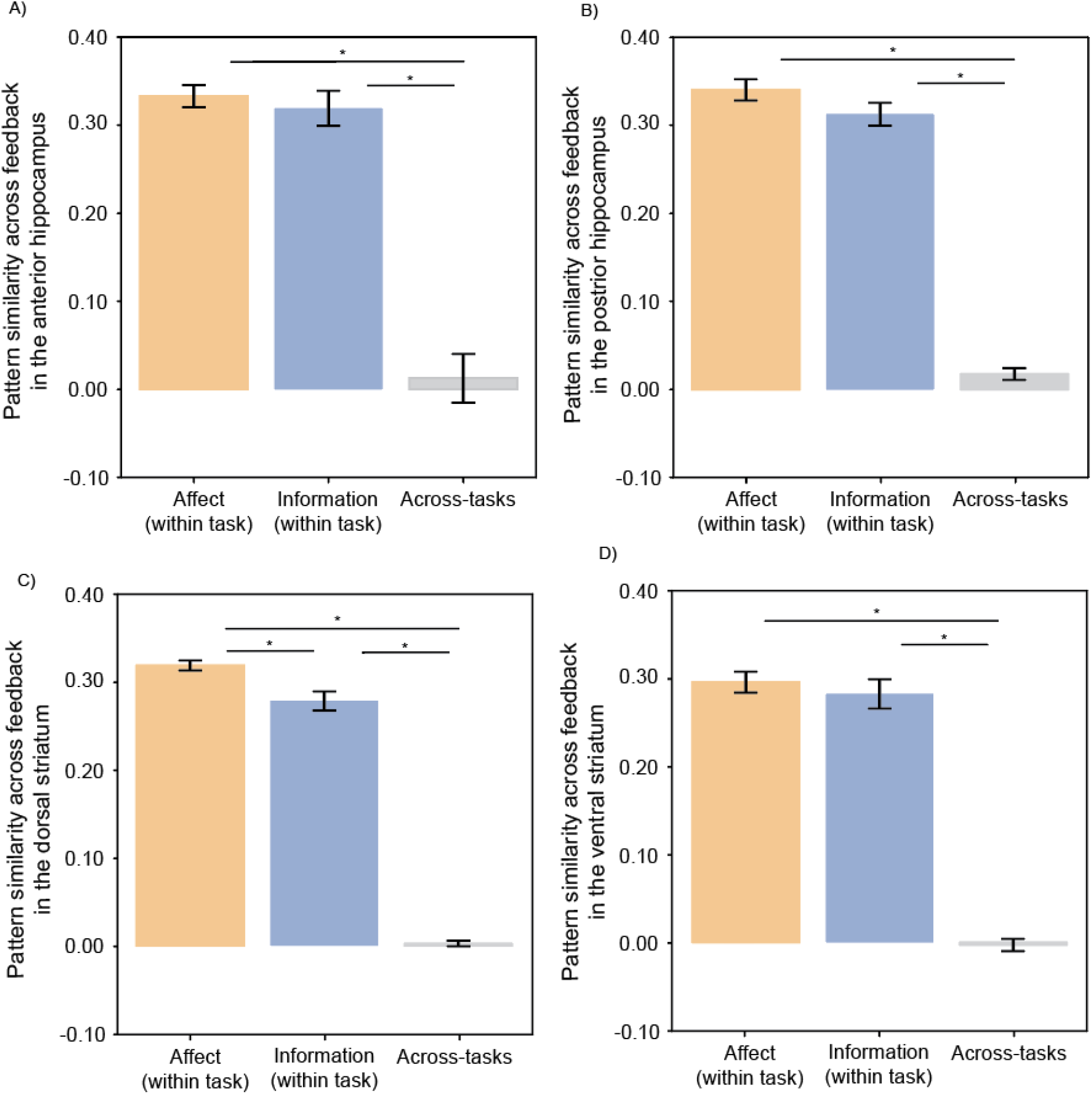
A) RSA effects in the anterior hippocampus. The anterior hippocampus showed greater pattern-similarity for within-task than across-task representation of affective and informative components of reward receipt. B) RSA effects in the posterior hippocampus. The posterior hippocampus showed greater pattern similarity for both the affective and informative card tasks. C) RSA effects in the dorsal striatum. The dorsal striatum showed greater pattern similarity for both the affective and informative card tasks. Moreover, the dorsal striatum showed greater pattern similarity for the affective components of reward receipt than the informative components of reward receipt. D) RSA effects in the ventral striatum. The ventral striatum showed greater pattern similarity for both the affective and informative card tasks.

### 3.3. Are the striatum and hippocampus sensitive to affective and informative reward magnitude?

Next, we tested whether the striatum and the hippocampus are sensitive to affective and informative reward magnitude. We constructed a hypothesized model, where we assumed pattern similarity increased linearly as reward magnitude increased. In this analysis, a positive correlation between the hypothesized model and the across-run pattern similarity in the affective card task indicates greater pattern similarity for high than low reward. A positive correlation between the hypothesized model and the across-run pattern similarity in the informative card task indicates greater pattern similarity for more common (predicted) than rare (unpredicted) feedback.

We did not find significant correlations between the hypothesized model and observed across-run pattern similarity in the anterior and posterior hippocampus (figure 4A and 4B) for both the affective (anterior hippocampus: t(18) = -0.49; *p* = 0.63; posterior hippocampus: t(18) = 0.74; *p* = 0.47) and informative (anterior hippocampus: t(18) = 0.02; *p* = 0.99; posterior hippocampus: t(18) = 0.97; *p* = 0.35) components of reward receipt. Direct comparisons between affective and informative components of reward also showed no significant difference in sensitivity to reward magnitudes in both the anterior (t(36) = -0.38; *p* = 0.70) and posterior hippocampus (t(36) = -0.23; *p* = 0.82).

**Figure 4.**
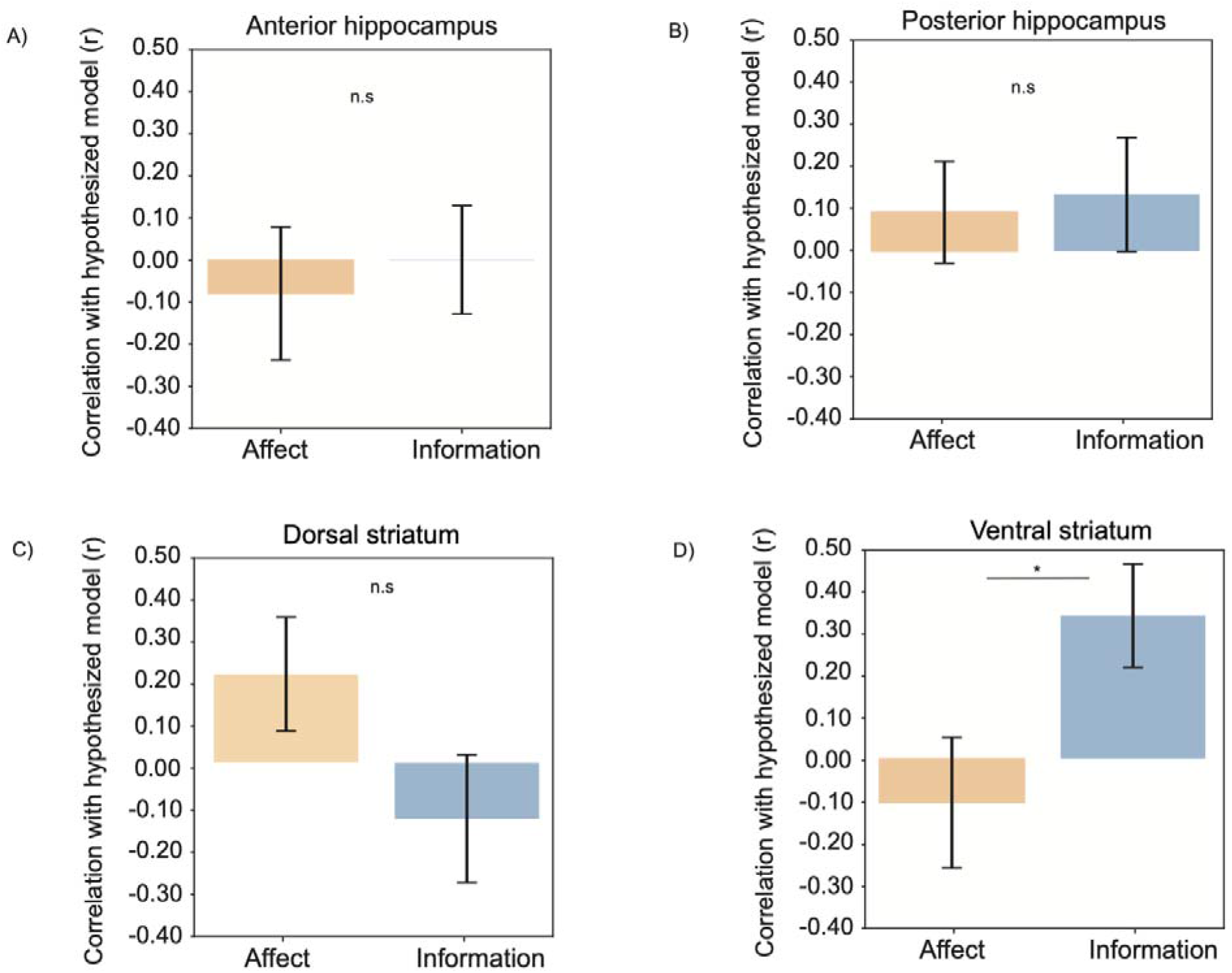
A & B & C) In the anterior and posterior hippocampus, and dorsal striatum, there was no correlation between the hypothesized model and across-run pattern similarity for both the affective and informative card tasks, suggesting that the anterior and the posterior hippocampus and the dorsal striatum were not sensitive to affective and informative reward magnitude. D) In the ventral striatum, there was a positive correlation between the hypothesized model and across-run pattern similarity for the informative card task, suggesting that the ventral striatum pattern similarity increased with the predictiveness of informative feedback. There was no correlation between the hypothesized model and across-run pattern similarity for the affective card task, suggesting a differential coding schema for affective and informative components of reward receipt in the ventral striatum.

We also did not find that significant correlation between the hypothesized model and the observed across-run pattern similarity for both the affective card task (t(18) = 1.66, *p* = 0.12) and the informative card task (t(18) = -1.28; *p* = 0.22) in the dorsal striatum (figure 4C). Direct comparison between the affective and information components of reward receipt also showed no significant difference in sensitivity to reward magnitude (t(36) = 1.65; *p* = 0.11), suggesting that the dorsal striatum was not sensitive to either affective or informative reward magnitude. Given that we have a relatively small sample size. Further research is needed to determine whether the observed null effect was due to insufficient statistical power or if the dorsal striatum was not sensitive to affective reward magnitude.

In contrast, we found that, in the ventral striatum (figure 4D), there was a positive correlation between the hypothesized model and the observed across-run pattern similarity for the informative components of reward (t(18) = 2.69, *p* = 0.02), such that ventral striatum showed increased pattern similarity with predictiveness of informative feedback. However, we did not find a significant correlation for the affective components of reward receipt (t(18) = - 0,65, *p* = 0.52), suggesting that the ventral striatum was only sensitive to the predictiveness of information. A direct comparison between affective and informative components of reward receipt showed a significant difference in sensitivity to reward magnitude (t(36) = -2.18, *p* = 0.04), suggesting that the ventral striatum has differential coding schema for affective and informative components of reward receipt, such that the ventral striatum is only sensitive to the predictiveness of information, but not to affective reward magnitude. Together, our findings suggested that only the ventral striatum showed differential sensitivity to affective and informative reward magnitudes, such that ventral striatum showed greater activation to more predictive informative feedback, but was not sensitive to affective reward magnitude.

## 4. Discussion

To understand how affective and informative components of reward receipt are represented in the striatum and the hippocampus, we tested participants on two card guessing tasks that separated affective and informative components of reward receipt. Using high-resolution fMRI and representation similarity analysis, we found that both the striatum and the hippocampus differentially represent affective and informative components of reward receipt, such that all four ROIs – anterior hippocampus, posterior hippocampus, dorsal striatum, and ventral striatum, showed higher pattern similarity within versus across tasks. Higher within-task pattern similarity indicates the neural representations were more consistent within the same task than between tasks. Moreover, we found that the dorsal striatum showed greater pattern similarity for affective card task than the informative card task, suggesting stronger representations of affect versus information in this region. There was no significant difference in the pattern similarity for affective and informative card tasks in the ventral striatum and the hippocampus. Further, we found that the ventral striatum was sensitive to the predictiveness of information, such that across-run pattern similarity increased with the more common information. Together, these findings suggest that affective and informative components of rewards might be represented differently in the striatum, though future studies are needed to investigate how affective and informative components of rewards are represented in the brain.

Our findings contribute to an expanding body of research linking striatal activation to the receipt of rewards (Delgado et al., 2000; Kahnt et al., 2009; Smith & Delgado, 2015). Consistent with prior literature, we showed that both the dorsal (Balleine et al., 2007; Delgado et al., 2000, 2003) and ventral striatum (Smith et al., 2016) are responsive to the affective components of reward receipt. We also found higher within-task than across-task pattern similarity for informative components of reward receipt in both dorsal and ventral striatum, which extends prior work by showing that the dorsal and ventral striatum are not only important for affective components of reward receipt, but are also important for informative components of reward receipt (Tricomi & Fiez, 2012).

Furthermore, consistent with our hypothesis that the hippocampus would also be important for informative feedback, we found both the anterior and posterior hippocampus showed greater pattern similarity for informative components of reward receipt. This finding aligns with previous research suggesting the involvement of the hippocampus in processing informative aspects of stimuli, demonstrating the importance of the hippocampus in integrating contextual information, and supporting its involvement in processing informative feedback during reward tasks (Charpentier et al., 2018; Cowan et al., 2021). Interestingly, we found that both the anterior and posterior hippocampus are important for affective information as well, providing support that the hippocampus also contributes to reward-related behaviors. Together, our findings contribute to a deeper understanding of the neural mechanisms underlying reward processing and provide more comprehensive insights into the representation of affective and informative components of reward receipt in the striatum and the hippocampus.

Previous literature has suggested there might be a dichotomy between the dorsal and ventral striatum in reward processing. The dorsal striatum has been often linked with processing informational aspects of rewards, while the ventral striatum is often associated with responding to affective cues (Filimon et al., 2020; O’Doherty et al., 2004; Tricomi & Fiez, 2012). We also found a distinction in the dorsal striatum and ventral striatum for processing affective and informative components of reward receipt. However, unlike previous work (Cardinal et al., 2002; Pool et al., 2022), our results showed a different pattern. We found that affective components of reward receipt were represented more similarly in the dorsal striatum, but not in the ventral striatum, which suggests that while the dorsal striatum is traditionally associated with processing informational aspects of rewards (Davidson et al., 2004; O’Doherty et al., 2004; Tricomi & Fiez, 2012; Vo et al., 2014), it is also responsiveness to affective cues and hedonic experiences, which advances our understanding of how distinct reward components are processed in the striatum. Future studies using our approach which combines high-resolution imaging and RSA in a task that manipulates information versus affect of anticipatory cues could help understand how these regions differentially respond to these two features of reward.

Furthermore, our study revealed that the ventral striatum was sensitive to the predictiveness of information, such that there was greater pattern similarity in response to more common feedback. This finding aligns with previous literature implicating the role of the ventral striatum in reward prediction and reinforcement learning processes (Daw et al., 2006; O’Doherty et al., 2004). We speculate that this observed pattern of activation reflects the ventral striatum’s crucial role in signaling the anticipation of forthcoming rewards based on the degree of predictability inherent in the provided information. Specifically, the ventral striatum is known to integrate various contextual cues and environmental stimuli to generate predictions about future rewards, facilitating adaptive decision-making and goal-directed behaviors (Filimon et al., 2020). The heightened ventral striatum activation observed in response to more predictive cues suggests that it plays an important role in encoding and processing the significance of incoming information. However, future studies with a larger sample size are needed to confirm the relationship. Together, our findings shed light on how the ventral striatum integrates predictive information to guide behavior.

Although our findings advance our understanding of how the striatum encodes different components of reward receipt, we note that our work was accompanied by two caveats. First, we did not have a direct measure of affective states during the affective card task and it is difficult to separate affective and informative components completely. Second, unlike our prior results for the informative card task (Smith et al., 2016), we found that participants were more likely to stay with a choice following expected (i.e., unsurprising) feedback. Although we speculate that this difference may due to the fact that all trials in the present study contained feedback, future studies are needed to understand how information value is shaped by motivational states. Future studies are needed to understand how information value and choice behaviors are shaped by motivational states. We speculate that the anticipation of unexpected outcomes may trigger a heightened motivational response, leading to a greater likelihood of information-seeking and choice-switching following surprising feedback (Shen et al., 2023). Taken together, future work using alternative tasks is needed to separate affective and informative components of reward and to better understand how the striatum and the hippocampus respond to different aspects of rewards upon receipt.

In summary, our work provided insights into the representation of affective and informative components of reward receipt in the striatum and the hippocampus. We showed evidence that both the striatum and the hippocampus are important to the affective and informative components of reward receipt. On top of that, we also found that the dorsal striatum showed greater pattern similarity to affective components of reward than informative components of reward and finally, we found the ventral striatum was sensitive to predictiveness of information, but not to affective reward magnitude, suggesting differential coding schema for affective and informative components of reward receipt in the striatum.

## Data and code availability

Analysis code related to this project can be found on GitHub: https://github.com/KarenShen21/ru-highres

## Author contribution

K.S.W., D.V.S., and M.R.D. designed research; K.S.W performed research; X.S., D.V.S and V.P.M. analyzed data; and X.S, D.V.S., and V.P.M wrote the paper.

## Competing Interest Statement

No conflict of interest.

## Notes

### Competing Interest Statement

The authors have declared no competing interest.

